# A3D Database: Structure-based Protein Aggregation Predictions for the Human Proteome

**DOI:** 10.1101/2021.11.17.468872

**Authors:** Aleksandra E. Badaczewska-Dawid, Javier Garcia-Pardo, Aleksander Kuriata, Jordi Pujols, Salvador Ventura, Sebastian Kmiecik

**Affiliations:** Department of Chemistry, Iowa State University, Ames, 50011 IA, USA; Biological and Chemical Research Center, Faculty of Chemistry, University of Warsaw, Pasteura 1, 02-093 Warsaw, Poland; Institut de Biotecnologia i de Biomedicina (IBB) and Departament de Bioquímica i Biologia Molecular, Universitat Autònoma de Barcelona, 08193 Bellaterra, Barcelona, Spain

## Abstract

**Motivation:** Protein aggregation is associated with highly debilitating human disorders and constitutes a major bottleneck for producing therapeutic proteins. Our knowledge of the human protein structures repertoire has dramatically increased with the recent development of the AlphaFold (AF) deep-learning method. This structural information can be used to understand better protein aggregation properties and the rational design of protein solubility. This article uses the Aggrescan3D (A3D) tool to compute the structure-based aggregation predictions for the human proteome and make the predictions available in a database form.

**Results:** Here, we present the A3D Database, in which we analyze the AF-predicted human protein structures (for over 17 thousand non-membrane proteins) in terms of their aggregation properties using the A3D tool. Each entry of the A3D Database provides a detailed analysis of the structure-based aggregation propensity computed with A3D. The A3D Database implements simple but useful graphical tools for visualizing and interpreting protein structure datasets. We discuss case studies illustrating how the database could be used to analyze physiologically relevant proteins. Furthermore, the database enables testing the influence of user-selected mutations on protein solubility and stability, all integrated into a user-friendly interface.

**Availability and implementation:** A3D Database is freely available at: http://biocomp.chem.uw.edu.pl/A3D2/hproteome

## 1 Introduction

In July 2021, a database of highly accurate structure predictions for the human proteome was published (Tunyasuvunakool *et al.*, 2021). The predictions were computed using the newly developed neural network-based model AlphaFold (AF). They were shown to be competitive with experimental structures and more accurate than those of alternative methods (Jumper *et al.*, 2021). The newly reported structural information allows generating proteome-wide repositories reporting on globular proteins’ aggregation properties.

Here we have constructed the AGGRESCAN3D (A3D) Database by computing the aggregation propensity of the human protein models from the AF database. The A3D is a structure-based predictor of protein aggregation developed by our group to identify and quantify surface-exposed aggregation-prone regions (S-APRs). The A3D algorithm exploits the information of three-dimensional atomic models to compute the structurally corrected aggregation values (A3D score) for each particular amino acid within the protein (Kuriata, Iglesias, Kurcinski, *et al.*, 2019; Kuriata, Iglesias, Pujols, *et al.*, 2019; Pujols *et al.*, 2018; Zambrano *et al.*, 2015). A3D outperforms classic sequence- and composition-based algorithms when dealing with globular proteins in their native conformations and can compute the effect of single or multiple user-defined mutations on protein stability and aggregation propensity, as well as automatically suggesting solubilizing amino acid changes. This algorithm has been employed to study the constraints imposed by aggregation on protein evolution (Carija *et al.*, 2019), to diagnose the functional impact of genetic mutations (Seaby and Ennis, 2020), to predict the aggregation of the SARS-CoV-2 proteome (Flores-León *et al.*, 2021), to assist the design of novel nanomaterials (Gil-Garcia and Ventura, 2021), or to engineer the solubility of therapeutic proteins (de Aguiar *et al.*, 2021; Gil-Garcia *et al.*, 2018) among many other applications.

The A3D database will be helpful in the study and redesign of human proteins’ solubility. It will also allow investigating correlations between structural aggregation propensity and protein function, stability, architecture, location, abundance, lifetime, or essentiality at the proteome level. In addition, it might constitute a valuable tool to decipher the aggregative properties of specific subproteomes associated with different diseases. We illustrate the performance and utility of the database with selected case reports.

## 2 Methods

### 2.1 Database construction

The AF Database collects 23391 protein structure predictions assigned to the *Homo sapiens* proteome, corresponding to 20504 unique UniProt entries (Jumper *et al.*, 2021). This set covers well the subset of human proteins with reviewed status, which are longer than 16 amino acids. The extremely long sequences (longer than 2700 residues and up to 34350) were split into overlapping fragments and modeled independently. As a result, the AF database provides multiple structure predictions, a few to several dozen, for a particular UniProt identifier. The membrane proteins have specific physicochemical properties on their surface, which can significantly bias the A3D predictions. Thus, we excluded a total of 5156 transmembrane and intramembrane proteins, which can be further analyzed with a custom configuration of A3D settings. For all the proteins, we collected the additional information’s that can be used in database searching. They include UniProt identifier, gene, common short name, and long descriptive name. For the user’s convenience, we added information about the length of the original sequence. In addition, for sequences split into fragments, we provided the region corresponding to the predicted structure. It can make it much easier to identify a domain of interest to the user. In total, the A3D database contains over 17 thousand structural deposits, corresponding to 15349 unique UniProt IDs (individual proteins, because for some cases, the original sequence was split into fragments). Because the predicted structures vary considerably in confidence level (pLDDT), we also report for each entry the number of residues below a threshold of 70 and 50, which can help to screen out underpredictions for user-defined A3D analysis. The unified and integrated metadata accompanied by referencing identifiers in the A3D database is available for download in CSV format from the Download tab of the A3D database webpage.

### 2.2 A3D analysis

We performed an A3D analysis for 17503 structures from the AF database. For each case, we tested the aggregation properties of three variants. The first was the original AF-predicted structure, while the second and third were modified structures deficient in residues with a pLDDT score lower than 70 and 50, respectively. We run all jobs through the RESTful service of Aggrescan3D 2.0 web server with default A3D settings, i.e., with 10 Å distance of aggregation prediction and FoldX-based energy minimization for stability calculations.

All results are freely available online in the A3D database http://biocomp.chem.uw.edu.pl/A3D2/hproteome

## 3 Results

### 3.1 A3D Database features

The A3D database integrates A3D predictions for over 17 thousand non-membrane proteins from the AF database. Membrane proteins were excluded because although A3D accurately predicts hydrophobic transmembrane regions as highly aggregation-prone, they are protected by the membrane lipidic bilayer and not exposed to the solvent. AF produces a per-residue estimate of each model’s confidence on a scale from 0 – 100 (Tunyasuvunakool *et al.*, 2021). This confidence score (pLDDT) reports on the quality of the AF prediction (Jumper *et al.*, 2021). To evaluate A3D predictions on AF-derived models, we randomly selected 100 protein entries and inspected them (Table S1). Manual data curation revealed that low pLDDT scores might result in misleading A3D predictions, because often they correspond to protein regions that are either more exposed or sheltered in the model than in their native/natural conformation. Therefore, after testing different pLDDT thresholds, we decided to precompute A3D on top of three different AF models for each protein entry: The full-length protein model and two additional models in which residues with pLDDT < 70 or residues with pLDDT < 50 were removed (see Supplementary Materials).

The A3D database is presented as the main A3D search front page. The content of the database can be queried by UniProt ID, Gene, or by protein name (see S1 Movie in Supplementary Materials for the short tutorial). By selecting the desired target in the results list, each protein entry can be accessed, leading to the A3D results pages. The A3D predictions are presented in a series of tabs that link to pages containing: (a) the project details, (b) an interactive A3D profile, (c) a detailed table containing A3D scores and AF pLDDTs at the residue level (d), the structural information, (e) customizable calculations and (f) an image gallery. In the *Structure* page, protein structures can easily be visualized and analyzed interactively. Two different models are presented for each entry (see Figure 1). The top model reports on the A3D score, while the bottom model depicts the AF pLDDTs. The *Custom Jobs* page contains links to the precalculated information for the entry at the two preselected AF confidence cutoffs (70 and 50), a residue editor, and the possibility to submit a new job to A3D with a custom-selected AF pLDDT cutoff (see Notes in Supplementary Materials). In addition, a mutation editor allows the introduction of one or multiple mutations and submits a new job to A3D, where the predicted changes in solubility and stability can be retrieved. New jobs will be immediately listed and accessible in the A3D queue.

**Figure 1.**
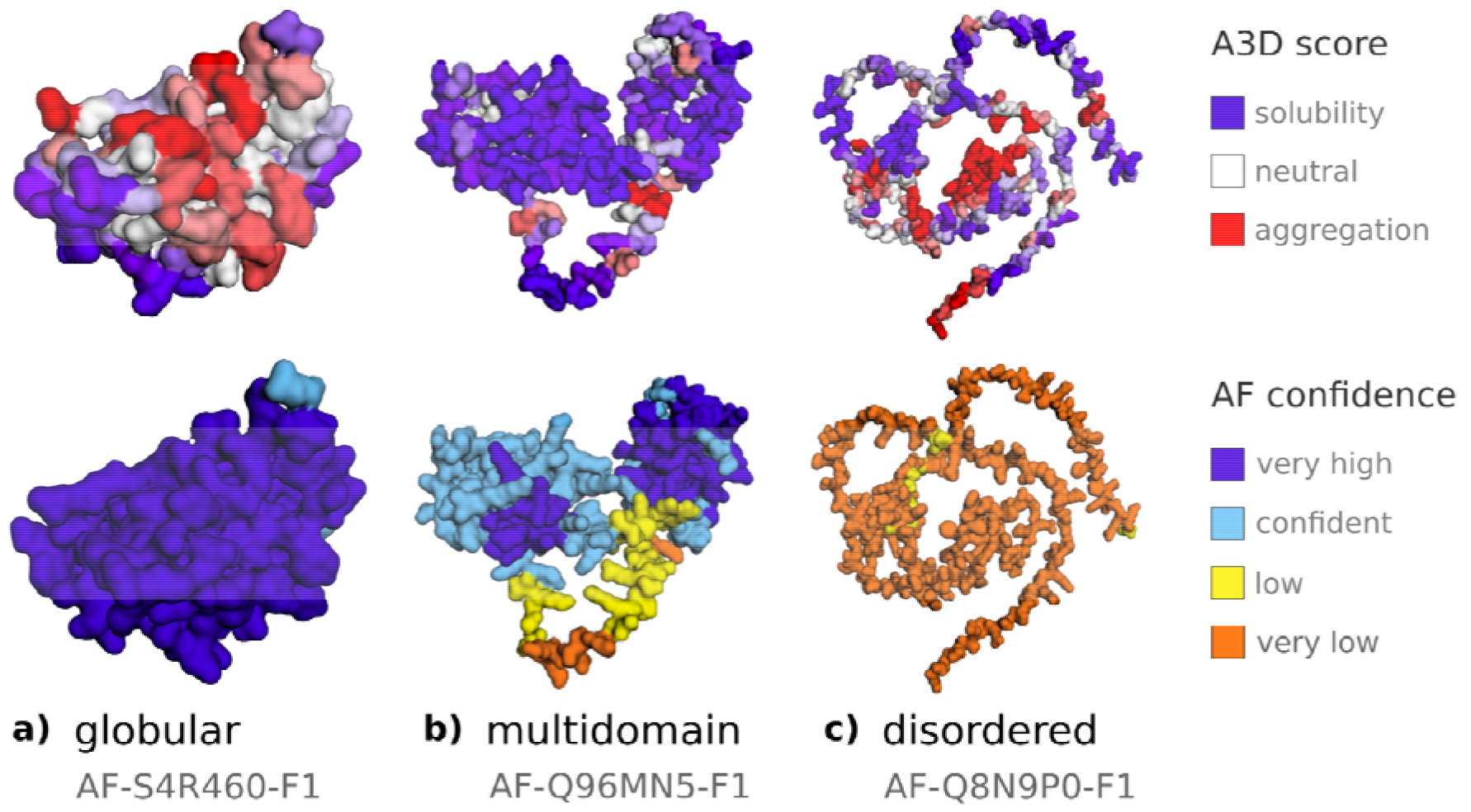
Examples of protein model visualizations from the A3D database. For each database entry, under the *Structure* tab, protein models are visualized in an interactive way. Two protein copies are presented colored according to (i) the A3D score and (ii) the AlphaFold (AF) model confidence score. The A3D score is visualized in shades from dark blue (highly soluble residues, score > +2.5), through white (no predicted influence on aggregation properties), to dark red (aggregation-prone residues, score < −2.5). The AF per-residue confidence score (pLDDT) is presented in dark blue (very high confidence, pLDDT > 90), light blue (confident, 90 > pLDDT > 70,), yellow (low confidence, 70 > pLDDT > 50), orange (very low confidence, pLDDT < 50). Note that pLDDT < 50 is a reasonably strong predictor of disorder (Tunyasuvunakool, *et al.*, 2021), which suggests that a particular region may be unstructured as a linker between domains (see panel b) or as an inherently disordered domain (see panel c). Panel a) shows an example of a globular protein predicted with high confidence.

### 3.2 Example cases

Below we investigate two example cases. In the first example, we analyzed the structure of the human Copper-Zinc Superoxide Dismutase (SOD1), for which a variety of mutations underly the formation of protein deposits in familial amyotrophic lateral sclerosis (Deng *et al.*, 1993). A comparative analysis of the structural aggregation propensity of this protein was done with A3D using either the experimental PDB coordinates or its equivalent AF-derived model (Figure 2). In both cases, A3D detected the presence of a strong S-APR that overlaps with the predicted dimerization interface (Figure 2), explaining why mutations favoring dissociation promote SOD1 deposits (Elam *et al.*, 2003).

**Figure 2.**
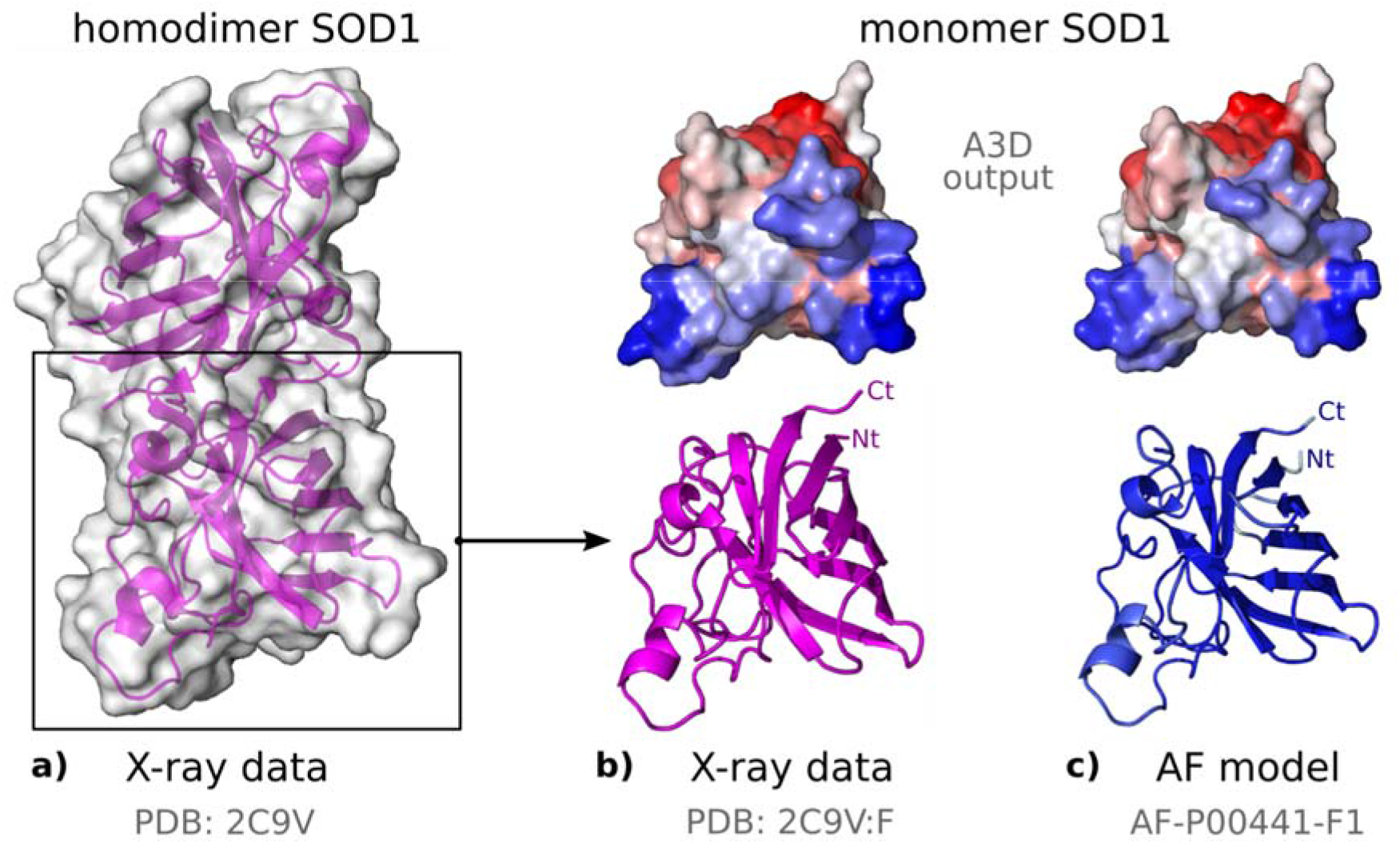
Comparison of the aggregation propensity of human Copper-Zinc Superoxide Dismutase (SOD1) calculated with A3D using PDB data and the equivalent AlphaFold2 model. a) Crystal structure of SOD1 homodimer (PDB ID: 2C9V (Strange et al. 2006)) b) SOD1 X-ray structure (PDB ID: 2C9V:F, bottom panel) and its A3D analysis (top panel). C) SOD1 AlphaFold model (AF-P00441-F1, bottom panel) and its A3D analysis (top panel). A3D analysis is presented using coloring scheme in a gradient from blue (high solubility) to white (no impact on protein aggregation) to red (high aggregation propensity).

In the second example, we compared human procarboxypeptidase A3 (hpCPA3) and human procarboxypeptidase B (hpCPB) (Figure 3). Both exopeptidases share a preference for basic residues, but whereas hpCPB is pancreatic, hpCPA3 is mast-cell-specific. In contrast to hpCPB, no experimental structure exists for hpCPA3. The AF-model for the hpCPA3 backbone is virtually identical to the one of the hpCPB crystal structure. However, the A3D analysis indicates a different S-APRs distribution, with hpCPA3 being clearly more soluble than hpCPB (Figure 3), with average A3D scores of −0.79 and −0.68, respectively. According to A3D, the higher solubility of hpCPA3 responds to the presence of a positively charged surface patch, absent in hpCPB, and likely necessary for the binding of the protein to proteoglycans within mast cell granules (Pejler *et al.*, 2009).

**Figure 3.**
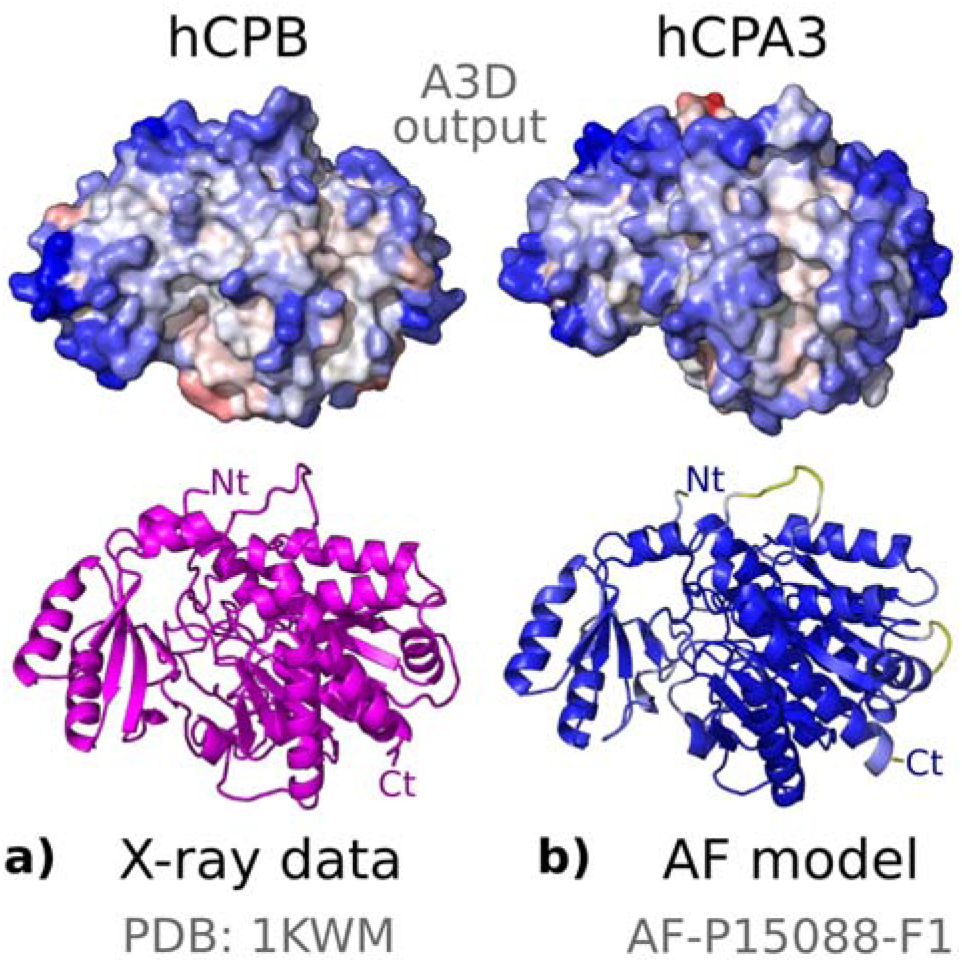
Aggregation propensity of human procarboxypeptidases B and A3 predicted with A3D. a) Structural model of human procarboxypeptidase B (hpCPB) (PDB code: 1KWM (Barbosa Pereira *et al.* 2002), residues 16-417) in a ribbon representation (bottom panel) and aggregation propensity of human hpCPB calculated with A3D (top panel). b) AlphaFold model of the human procarboxypeptidase A3 (hpCPA3) (or Mast Cell Carboxypeptidase, AlphaFold2 code: AF-P15088-F1, residues 16-417) in a ribbon representation (bottom panel) and A3D analysis of human hpCPA3 (top panel). The protein surface is colored according to the A3D score.

### 3.3 Prediction of the impact of mutations on protein solubility and stability

A3D has been shown to be accurate in predicting the changes in the solubility of globular proteins upon mutation (Zambrano *et al.*, 2015). We compared the performance of A3D when modeling the impact of mutations on the solubility on top of either the experimental structures and the equivalent AF-derived models, using the mutation editor in the A3D database. To this aim, we selected a reduced set of structurally and sequentially unrelated proteins (Table S2 and Figure 4). The predictions turned out to be accurate and coincident for all the proteins, independently of the kind of structural input we used. In addition to changes in solubility, A3D provides the impact of the selected mutations on protein stability, as calculated by FoldX (Schymkowitz *et al.*, 2005). Importantly, it has been shown that the quality of AF models is sufficient to predict protein solubility changes upon mutation using FoldX(Akdel *et al.*, 2021).

**Figure 4.**
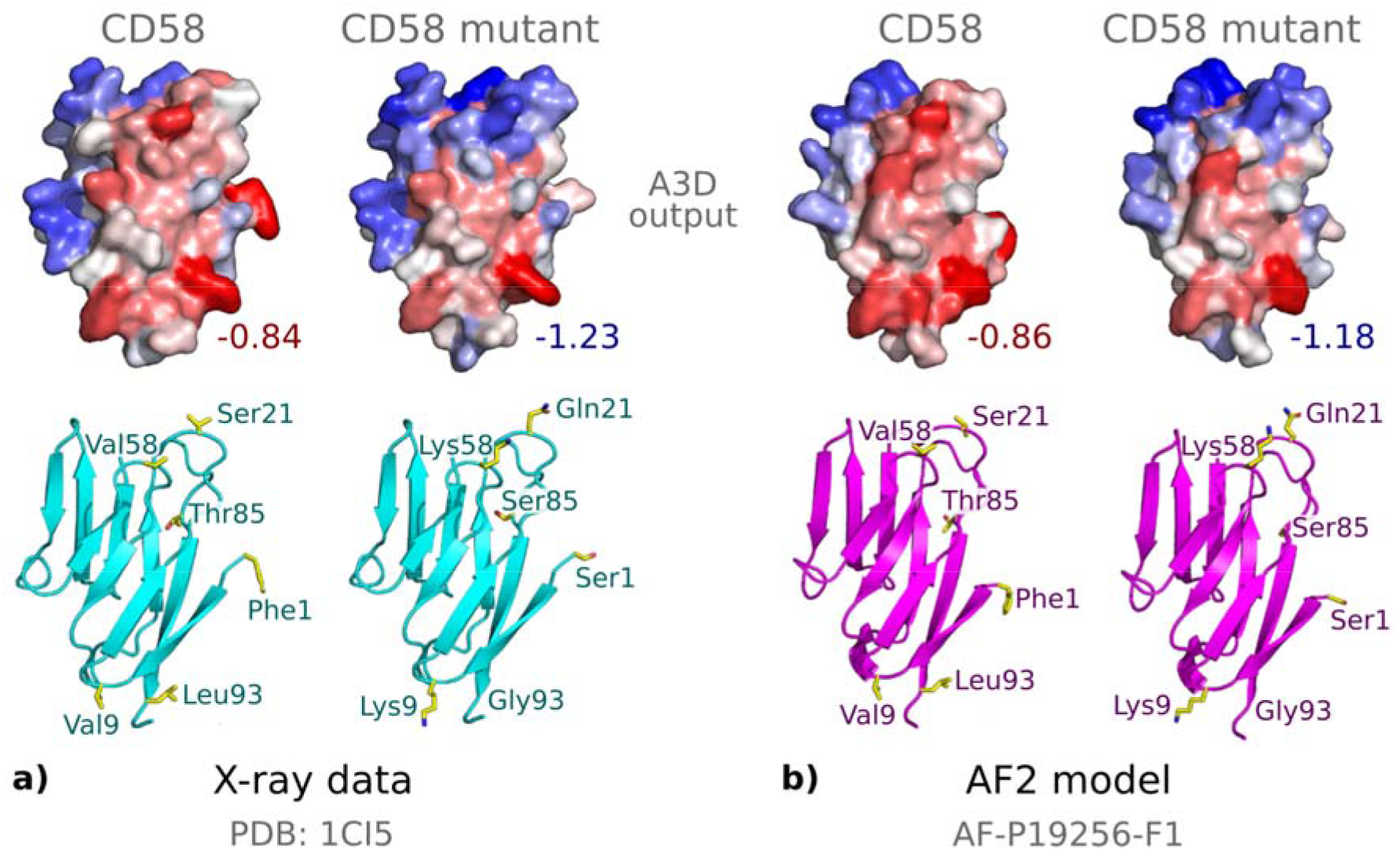
Comparison of the performance of A3D in predicting the changes in the solubility of Lymphocyte function-associated antigen 3 (CD58) between a) the experimentally determined structure (PDB ID: 1CI5 (Sun *et al.*, 1999), colored in cyan) and b) the Alphafold2-derived model (AF-P19256-F1, colored in magenta). The top row in both panels shows A3D analysis for the wild-type CD58 (WT) and its solubility-optimized mutant (see Table S1 for details), respectively. For both sources of structural data, the introduced mutations (F1S, V9K, S21Q, V58K, T85S, L93G) result in a more soluble CD58 variant. The calculated average A3D scores improve from −0.84 to −1.23 for X-ray structure, and from −0.86 to −1.18 for the AF2 model. In all the cases, surface representations are colored according to the predicted A3D score, as defined in Figure 2.

## 4 Discussion

A3D is a structure-based algorithm that uses three-dimensional protein coordinates to project each amino acid’s intrinsic aggregation propensity value into the structure. Then, the aggregation score is corrected as a function of its specific solvent exposure and the aggregation propensity of neighboring residues within a 10 Å sphere of radius. Given this dependence on the spatial position of each atom of the protein, both the atomic resolution and the biological relevance of the input structures impact A3D prediction’s accuracy. Therefore, in order to correctly interpret A3D database aggregation tendencies, users might consider two critical characteristics of the AF database: (i) structure confidence and disorder content and (ii) quaternary structure context.

### 4.1 Interpreting structure confidence and disorder content

AF predicted structures cover 98.5% of the human proteome and involve complete protein chains in their monomeric state. Each amino acid in an AF model has a confidence score (pLDDT) that ranges from 0 to 100, indicating which regions could be considered equivalent to an experimentally determined structure. It is estimated that around 30% of the amino acids of the human proteome are located outside globular (well-folded) domains and constitute intrinsically disordered or low-complexity regions. Therefore, it is not surprising that AF-predicted structures frequently contain regions with very low (pLDDT < 50) scores (Ruff and Pappu, 2021). These permanently or transiently disordered segments are often not defined in experimental structures, and therefore were excluded from the A3D analysis. Now they are incorporated in the A3D Database. The A3D predictions are blind to the dynamic nature of these regions since AF models correspond to static single frames of the conformational ensemble. Therefore, the presence of disordered regions with low AF confidence might artificially shelter or increase the presence of S-APRs in adjacent high confidence protein domains.

To deal with this issue, the A3D dataset includes two additional precalculated structural aggregation profiles, which correspond to predicted structures with the pLDDT thresholds: pLDDT > 70 and pLDDT > 50 confidence, which was defined after manually curating the 100 AF models depicted in Table S1.

In the following points, we outline different situations users may face:

⍰ Proteins with high confidence values: the overall AF-predicted structure possesses high pLDDTs. All three computed structures converge into a unique aggregation prediction. Well-defined S-APRs can be delineated from A3D output structures (representative examples: Q68D91, S4R460, see panel a) in Figure 5).
⍰ Proteins with limited low confidence regions: proteins that display local surface modifications with confidence thresholds of >70% or >50% pLDDT. An increase or decrease of localized S-APRs can be observed (representative examples: P09382, Q9UKL6, Q5EE01, see panel d) in Figure 5).
⍰ Multidomain proteins: proteins with two or more globular domains that correspond to well-defined and confident regions, while tethering elements that connect globular domains possess low pLDDT values. The analysis with restricted pLDDT thresholds often results in disconnected domains and, as a consequence, some previously sheltered APRs might become exposed. As a result, S-APRs might diverge in the tree models, and their boundaries are less evident (representative examples: Q96MN5, Q96DA0, see panel b) in Figure 5). Note that for each database entry, the AlphaFold database provides Predicted Aligned Error (PAE) analysis. The PAE is useful for assessing whether relative domain positions are predicted correctly or not. If not, A3D analysis may be affected by incorrectly exposed domains.
⍰ Proteins with extended low-confidence regions: proteins with large low confidence regions localized in flexible C-, N-terminus, or long loops, usually displayed around a well-defined globular core. With the 70% and 50% pLDDTs cutoffs, disordered regions are often excluded from the model and, therefore, not considered in A3D prediction. As a result, the A3D predictions correspond to well-folded globular domains. Note that we might observe the presence of free amino acids in some cases, unattached to the protein main chain, essentially because they are in the exclusion limit between two pLDDT thresholds (representative examples: Q9NRE2, Q8NFW1, see panel e) in Figure 5).
⍰ Proteins with overall low confidence: usually, short polypeptides that are mostly disordered, exposing most of its surface to solvent. In some cases, we lose all the structure in predictions corresponding to >70% or >50% pLDDT threshold. (representative examples: Q8N9P0, Q8TEV8, Q9GZY1, see panels c) and f) in Figure 5).

**Figure 5.**
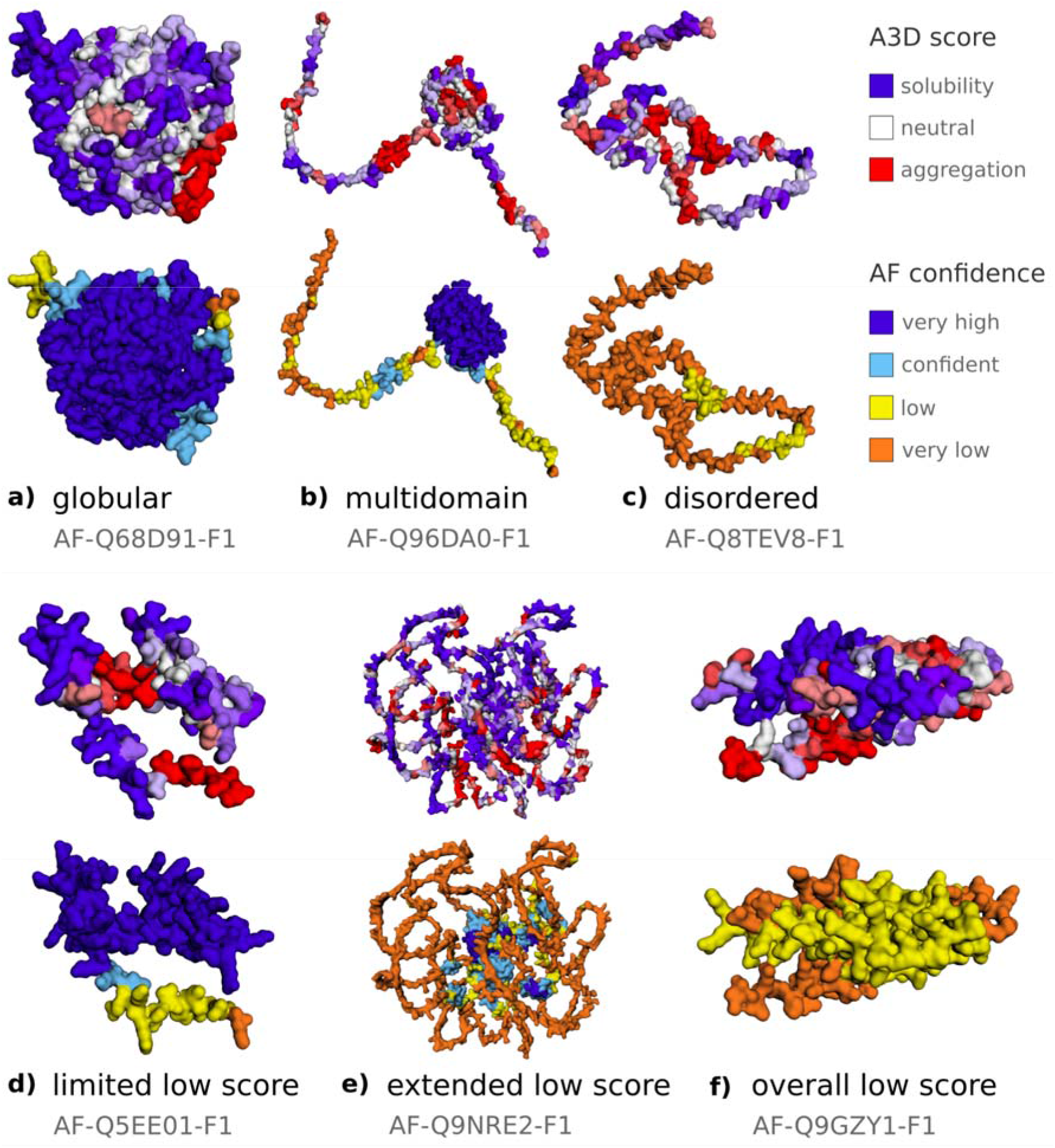
An overview of the different types of cases that may be encountered in the A3D database. The presented cases are colored using A3D score (top panels) and AF confidence score (bottom panels) and discussed in the 4.1 section.

### 4.2 Interpreting quaternary structure context

In the AF database, structure predictions are restricted to single chains. This precludes the analysis of the quaternary structure context in the A3D Database. The overlap between the physicochemical properties governing protein-protein native interactions and non-native contacts triggering aberrant self-assembly implies that protein interfaces are enriched in S-APRs. In proteins displaying quaternary structure, this results in an over-prediction of S-APRs in protein monomers, relative to native the same subunits in the oligomeric state.

### 4.3 Gathering important information

In this work, we suggest a general guideline to manage the A3D output. In addition, we strongly encourage users to compile as much information about their case study protein to exploit A3D precalculated predictions. Some helpful questions to shape the output would be:

i. Does the protein possess disordered regions?
ii. Do these disordered segments correlate with low pLDDT?
iii. Does the protein bind another protein? Does it form a homo- or hetero-oligomer?
iv. Does the protein contain a signal peptide?
v. Is the protein the mature form or on the contrary, is a proprotein?

In addition to the aforementioned pieces of advice, we also recommend reviewing dedicated bibliography on how to manage the A3D algorithm and how to evaluate its aggregation predictions (Pujols *et al.*, 2018; Kuriata, Iglesias, Pujols, 2019; Zambrano *et al.*, 2015; Santos *et al.*, 2020).

## 5 Conclusions

The A3D database makes available structural aggregation predictions at the residue level for over 17 thousand structural deposits with different AF model confidence intervals. When a high confidence AF model is available, the A3D predictions coincide with those made on top of the experimental structure. Thus, the database should allow assessment of the aggregation propensity of the human proteome and, eventually, engineering it in terms of solubility and stability.

## Acknowledgments

A.E.B-D. acknowledges financial support by Roy J. Carver Charitable Trust through Iowa State University Bioscience Innovation Postdoctoral Fellowship. This work was funded by the Spanish Ministry of Science and Innovation (MICINN) PID2019-105017RB-I00 to S.V, by ICREA, ICREA-Academia 2019 and by EU (PhasAge /H2020-WIDESPREAD-2020-5) to SV. J.G.-P. was supported by the Spanish Ministry of Science and Innovation with a Juan de la Cierva Incorporacion IJC2019-041039-I. SK acknowledges funding by the National Science Centre, Poland [2020/39/B/NZ2/01301].

## References

de Aguiar,R.B. et al. (2021) Generation and functional characterization of a single-chain variable fragment (scFv) of the anti-FGF2 3F12E7 monoclonal antibody. Sci. Rep., 11, 1432.

Akdel, M. et al. (2021) A structural biology community assessment of AlphaFold 2 applications. bioRxiv.

Barbosa Pereira, P.J., et al. Human procarboxypeptidase B: three-dimensional structure and implications for thrombin-activatable fibrinolysis inhibitor (TAFI). Journal of molecular biology 2002;321(3):537–547

Carija, A. et al. (2019) Computational Assessment of Bacterial Protein Structures Indicates a Selection Against Aggregation. Cells, 8.

Deng,H.-X. et al. (1993) Amyotrophic Lateral Ssclerosis and Structural Defects in Cu,Zn Superoxide Dismutase. Science, 261, 1047–1051.

Elam,J.S. et al. (2003) Amyloid-like filaments and water-filled nanotubes formed by SOD1 mutant proteins linked to familial ALS. Nat. Struct. Biol., 10, 461–467.

Flores-León,M. et al. (2021) In silico analysis of the aggregation propensity of the SARS-CoV-2 proteome: Insight into possible cellular pathologies. Biochim. Biophys. Acta: Proteins Proteomics, 1869, 140693.

Gil-Garcia,M. et al. (2018) Combining Structural Aggregation Propensity and Stability Predictions To Redesign Protein Solubility. Mol. Pharm., 15, 3846–3859.

Gil-Garcia,M. and Ventura,S. (2021) Multifunctional antibody-conjugated coiled-coil protein nanoparticles for selective cell targeting. Acta Biomater., 131, 472–482.

Jumper,J. et al. (2021) Highly accurate protein structure prediction with AlphaFold. Nature, 596, 583–589.

Kuriata,A., Iglesias,V., Pujols,J., et al. (2019) Aggrescan3D (A3D) 2.0: prediction and engineering of protein solubility. Nucleic Acids Res., 47, W300–W307.

Kuriata,A., Iglesias,V., Kurcinski,M., et al. (2019) Aggrescan3D standalone package for structure-based prediction of protein aggregation properties. Bioinformatics, 35, 3834–3835.

Pejler,G. et al. (2009) Novel insights into the biological function of mast cell carboxypeptidase A. Trends Immunol., 30, 401–408.

Pujols,J. et al. (2018) AGGRESCAN3D: Toward the Prediction of the Aggregation Propensities of Protein Structures. Methods Mol. Biol., 1762, 427–443.

Ruff,K.M. and Pappu,R.V. (2021) AlphaFold and Implications for Intrinsically Disordered Proteins. J. Mol. Biol., 433, 167208.

Santos,J. et al. (2020) Computational prediction of protein aggregation: Advances in proteomics, conformation-specific algorithms and biotechnological applications. Comput. Struct. Biotechnol. J., 18, 1403–1413.

Schymkowitz,J. et al. (2005) The FoldX web server: an online force field. Nucleic Acids Res., 33, W382–8.

Seaby,E.G. and Ennis,S. (2020) Challenges in the diagnosis and discovery of rare genetic disorders using contemporary sequencing technologies. Brief. Funct. Genomics, 19, 243–258.

Strange, R.W., et al. Variable metallation of human superoxide dismutase: atomic resolution crystal structures of Cu-Zn, Zn-Zn and as-isolated wild-type enzymes. Journal of molecular biology 2006;356(5):1152–1162.

Sun, Z.Y., et al. Functional glycan-free adhesion domain of human cell surface receptor CD58: design, production and NMR studies. The EMBO journal 1999;18(11):2941–2949.

Tunyasuvunakool,K. et al. (2021) Highly accurate protein structure prediction for the human proteome. Nature, 596, 590–596.

Zambrano,R. et al. (2015) AGGRESCAN3D (A3D): server for prediction of aggregation properties of protein structures. Nucleic Acids Res., 43, W306–13.

