## Supplemental Information for "A3D Database: Structure-based Protein Aggregation Predictions for the Human Proteome"

**SUPPLEMENTARY INFORMATION**

**Movie S1.** A short tutorial on using the Aggrescan3D (A3D) database.

**Performance of A3D for the prediction of mutations using AlphaFold-derived models**

We used A3D to evaluate the impact of selected mutations on protein solubility. For each protein presented in Table S1, the changes in solubility upon mutation and expression have been previously characterized elsewhere [1]-[6]. The experimental data were compared with the result from A3D predictions using either PDB data or AlphaFold models.

**Table S1.** Randomly selected case study AF structures, grouped alphabetically by their Uniprot identifier.

A0A0B4J263, A0A1B0GVH4, A0A7I2V3D3, A6NC57, A6NKH3, A8MZ26, J3KSC0, O14602, O15091, O43278, O60258, O75044, O75626, O94901, O95613, P00748, P02776, P06400, P09382, P0CJ87, P10243, P13611, P16989, P20810, P23458, P28360, P34896, P40617, P46779, P49792, P52566, P55884, P60842, P63173, P84550, Q02224, Q05639, Q0VDD8, Q13151, Q13753, Q14533, Q15149, Q15831, Q17RW2, Q3LHN1, Q4UJ75, Q5EE01, Q5SVQ8, Q5TBA9, Q5VUJ9, Q68D91, Q6MZM9, Q6PF06, Q6UXY1, Q6ZRI0, Q709C8, Q7Z2R9, Q7Z6R9, Q86UW9, Q86YS6, Q8IWC1, Q8IYW2, Q8N2C9, Q8N660, Q8N9P0, Q8NDA2, Q8NFW1, Q8TC99, Q8TEV8, Q8WWM1, Q8WZ42, Q92626, Q969T7, Q96DA0, Q96GQ7, Q96K80, Q96MN5, Q96Q35, Q99218, Q99973, Q9BSV6, Q9BWX1, Q9BZ19, Q9GZY1, Q9H2X6, Q9H772, Q9HB66, Q9NPE2, Q9NRE2, Q9NV70, Q9NYC9, Q9P1Y6, Q9UBH0, Q9UHL9, Q9UKL6, Q9UND3, Q9Y243, Q9Y3F1, Q9Y5P4 and S4R460

**Table S2**. Comparison of the performance of AGGRESCAN3D to predict changes in protein solubility using experimentally determined structures and their equivalent AlphaFold2- derived models.

|  |  |  | **AGGRESCAN 3D PREDICTION** | | | |  |
| --- | --- | --- | --- | --- | --- | --- | --- |
| **Protein Name** | **Mutation/s** | **Solubility** | **PDB code** | **Result** | **AF code** | **Result** | **Ref** |
| Hemoglobin subunit beta | Glu6Leu | Decreased | 2D60 (B) | TN | AF-P68871-F1 | TN | [1] |
|  | Glu6Phe | Decreased | 2D60 (B) | TN | AF-P68871-F1 | TN | [1] |
|  | Glu6Trp | Decreased | 2D60 (B) | TN | AF-P68871-F1 | TN | [1] |
| CD58 | Phe1Ser  Val9Lys  Val21Gln  Val58Lys  Thr85Ser  Leu93Gly | Increased | 1CI5 (A) | TP | AF-P19256-F1 | TP | [2] |
| Translation initiation factor eIF2a | Ala27Gln  Leu46His  Val71Lys | Increased | 1Q8K (A) | TP | AF-P05198-F1 | TP | [3] |
| Interleukin 1 Beta | Leu10Asn | Decreased | 9ILB (A) | TN | AF-P01584-F1 | TN | [4] |
|  | Leu10Asp | Decreased | 9ILB (A) | TN | AF-P01584-F1 | TN | [4] |
|  | Lys97Gly | Decreased | 9ILB (A) | TN | AF-P01584-F1 | TN | [4] |
|  | Lys97Val | Decreased | 9ILB (A) | TN | AF-P01584-F1 | TN | [4] |
| Apolipoprotein D | Trp99His  Ile118Ser  Leu120Ser | Increased | 2HZR (A) | TP | AF-P05090-F1 | TP | [5] |
| Leptin | Trp100Glu | Increased | 1AX8 (A) | TP | AF-P41159-F1 | TP | [6] |
|  | His97Ser  Trp100Gln  Ala101Thr  Gly112Glu  Met136Ile  Trp138Gln  Gly145Glu | Increased | 1AX8 (A) | TP | AF-P41159-F1 | TP | [6] |
|  | Trp100Gln  Trp138Gln | Increased | 1AX8 (A) | TP | AF-P41159-F1 | TP | [6] |
| Abbreviations: TP, True positives; TN, true negatives; PDB code, Protein Data Bank accession number; AF code, Alpha Fold Protein Structure Database accession number. | | | | | | | |

**SUPPLEMENTARY REFERENCES**

**[1]** Adachi, K., et al. Effects of beta 6 aromatic amino acids on polymerization and solubility of recombinant hemoglobins made in yeast. The Journal of biological chemistry 1993;268(29):21650-21656.

**[2]** Sun, Z.Y., et al. Functional glycan-free adhesion domain of human cell surface receptor CD58: design, production and NMR studies. The EMBO journal 1999;18(11):2941-2949.

**[3]** Ito, T. and Wagner, G. Using codon optimization, chaperone co-expression, and rational mutagenesis for production and NMR assignments of human eIF2 alpha. Journal of biomolecular NMR 2004;28(4):357-367.

**[4]** Sim, J. and Sim, T. Amino acid substitutions affecting protein solubility: high level expression of streptomyces clavuligerus isopenicillin N synthase in Escherichia coli. Journal of Molecular Catalysis B: Enzymatic. 1999;6(3).

**[5]** Nasreen, A., et al. Solubility engineering and crystallization of human apolipoprotein D. Protein science: a publication of the Protein Society 2006;15(1):190-199.

**[6]** Ricci, M., et al. Misbehaving Proteins: Protein (Mis)folding, aggregation, and stability. New York; 2006.
